# Actin capping protein regulates actomyosin contractility to maintain germline architecture in *C. elegans*

**DOI:** 10.1101/2021.10.03.462551

**Authors:** Shinjini Ray, Priti Agarwal, Ronen Zaidel-Bar

## Abstract

Actin dynamics play an important role in the morphogenesis of cells and tissues, yet the control of actin filament growth takes place at the molecular level. A challenge in the field is to link the molecular function of actin regulators with their physiological function. Here, we report the *in vivo* role of the actin capping protein CAP-1 in the *C. elegans* germline. We show that CAP-1 is associated with actomyosin structures in the cortex and rachis, where it keeps the level of contractility in check. A 60% reduction in the level of CAP-1 leads to a 2-fold increase in F-actin and non-muscle myosin II and only a 30% increase in Arp2/3. CAP-1 depletion leads to severe structural defects in the syncytial germline and oocytes, which can be rescued by reducing myosin activity. Thus, we uncover a physiological role for actin capping protein in maintaining *C. elegans* fertility by regulating the level of actomyosin contractility.

## Introduction

Diverse actin structures support the shape and power the movement of biological entities from the subcellular to the tissue level. Their functional diversity, manifested by different actin network architectures and dynamics, is regulated by their association with a variety of actin binding proteins in a context-dependent manner (Pollard, 2016; Winder and Ayscough, 2005). The barbed end of the actin filament is a hotspot for regulation, where interactions between multiple proteins determines whether it will elongate or remain stable (Shekhar et al., 2016). Capping protein (CP, also known as CAPZ and β-actinin) is one of the major determinants of barbed end dynamics (Edwards et al., 2014). CP binding at the barbed end of the actin filament prevents addition or loss of actin subunits. It functions as a heterodimer composed of two structurally similar α and β subunits (Yamashita et al., 2003).

Cooperative and competitive interactions between CP, nucleators and elongation factors influence the actin network architecture, dynamics and mechanical properties. CP competes with formin, a nucleation and elongation factor, for the barbed end, where they transiently form a ternary ‘decision complex’. Binding of formin to a CP-bound barbed end weakens the association of CP and promotes elongation of the filament (Bombardier et al., 2015; Kim et al., 2010; Shekhar et al., 2015). Similarly, CP is in direct competition with the Ena/VASP family of actin elongation factors (Bear and Gertler, 2009). In a branched actin network, nucleated by the Arp2/3 complex, CP determines the length and thus indirectly controls the density of branching (Funk et al., 2021; Iwasa and Mullins, 2007; Pollard and Cooper, 2009). While Arp2/3 nucleates branches along the actin filaments, CP caps them. Shorter branches provide strength but have less flexibility (Akin and Mullins, 2008; Schaub et al., 2007; Vinzenz et al., 2012). Most of the studies revealing the function of CP at the molecular and subcellular levels were performed with *in-vitro* reconstitution assays or in cell culture systems (Blanchoin et al., 2000; Pernier et al., 2016; Shekhar and Carlier, 2017); the importance of capping protein at the tissue level is not yet fully characterized.

Force generated by actomyosin machinery is essential for cell and tissue morphogenesis as well as the physiological function of many tissues (Agarwal and Zaidel-Bar, 2019). One example is the syncytial germline of *C. elegans*, where inward contraction of a corset-like actomyosin structure lining the rachis balances the outward pulling force of germ cell membrane tension (Priti et. al, 2018). Whether CP contributes to actin architecture and indirectly affects actomyosin contractility has not yet been investigated in an *in-vivo* system. Here, we explored the role of CP in the nematode *C. elegans* and discovered that it is critically important for maintaining a balanced level of actomyosin contractility in the germline, and is therefore required for the proper structure and function of the tissue.

## Results

### CAP-1 is a component of the actomyosin corset in the syncytial germline

To explore the expression pattern and sub-cellular localization of CAP-1 in the adult *C. elegans* hermaphrodite, we generated a strain in which CAP-1 is endogenously tagged with the fluorescent protein mKate2 at its N-terminus. Imaging by spinning disk confocal microscopy revealed that CAP-1 is expressed in multiple adult tissues, including the germline, spermatheca, pharynx, and vulva (Figure 1A). Co-staining of mKate2::CAP-1 worms with the F-actin marker phalloidin showed that CAP-1 localizes exclusively to structures containing F-actin, and most, but not all, F-actin structures contain CAP-1 (Supplementary figure 1A). CAP-1 was observed to localize at the apical side of gut epithelial cells, facing the intestinal lumen, and it was found in striations along body-wall muscle cells and in the germline (Supplementary figure 1B). The *C. elegans* gonad, with two symmetrical U-shaped arms, contains a syncytium of germ cells, which first form sperm that is stored in the spermatheca and later form oocytes (Kimble and Crittenden, 2007). The germ cells exist as a syncytium, connected to a common germplasm-filled core called rachis via cytoskeletal rings known as rachis bridges (Bauer et al., 2021). In the germline, CAP-1 was expressed in all germ cells, where it localized to the cell cortices and surrounded the nuclei, and in the germline syncytium CAP-1 was enriched at rachis bridges (Figure 1B). CAP-1 was also observed in the cortices of early embryos, the significance of which will be explored elsewhere. Here, we have focused on the function of CAP-1 in the germline.

**Figure 1:**
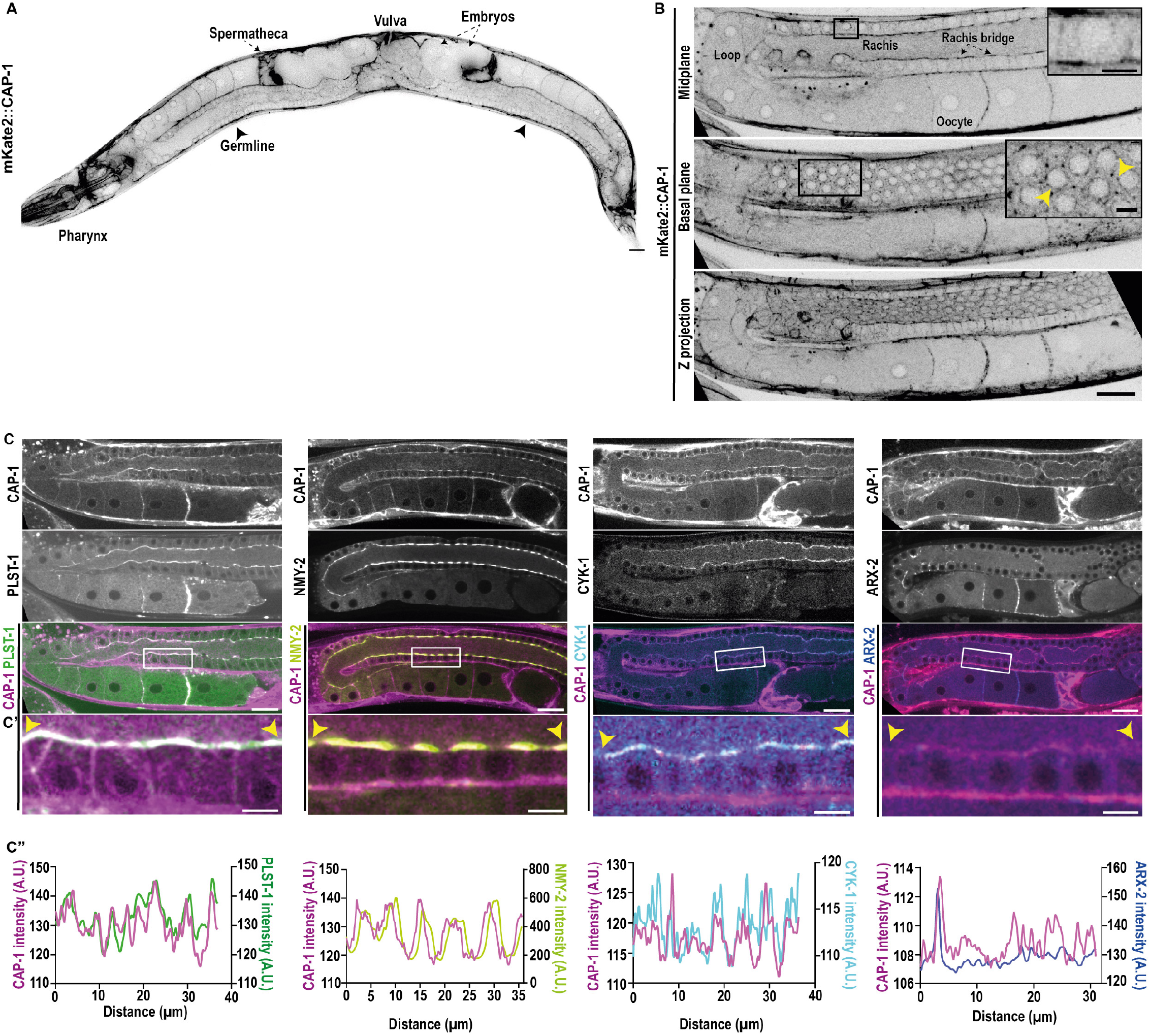
CAP-1 localizes at actomyosin-enriched structures in the *C. elegans* germline. (A) Inverted grayscale confocal fluorescence image of an adult *C. elegans* hermaphrodite expressing CAP-1endogenously tagged with mKate2. Arrows point to spermatheca and embryos. Arrowheads indicate germline. Scale bar, 20µm. (B) Representative inverted grayscale confocal fluorescence images of the *C. elegans* germline expressing mKate2::CAP-1. Midplane view showing enrichment of CAP-1 at rachis bridges and nuclei of germ cells, basal plane view showing localization at the germ cell cortex (arrowheads) and a maximum intensity Z-projection view showing CAP-1-enriched actomyosin corset in the gonad. Scale bar, 20µm. Inset scale bar, 5 µm. (C) Midplane view of the *C. elegans* germline showing colocalization of mKate2::CAP-1 with PLST-1::GFP, NMY-2::GFP, CYK-1::GFP and ARX-2::GFP. Scale bar, 20µm. (C’) Magnified view of the rachis region marked by a white rectangle in panel C. Scale bar, 5µm. (C’’) Fluorescence intensity profiles of endogenously-tagged proteins along a 5-pixel wide line drawn along the rachis between the arrowheads marked in panel C’.

We have previously shown that the syncytial germline architecture is maintained by contractility of an actomyosin corset surrounding the rachis (Priti et. al. 2018). To further investigate the localization of CAP-1 with respect to other components of the actomyosin corset, we crossed the mKate2::CAP-1 strain with strains expressing endogenously GFP-tagged PLST-1/plastin, NMY-2/non-muscle myosin II, CYK-1/diaphanous formin, and ARX-2/Arp2. As evident from the images and corresponding intensity line profiles, CAP-1 nearly perfectly co-localized with the actin cross-linker PLST-1 and NMY-2 along the rachis (Figure 1C, C’, C’’). CAP-1 and the formin CYK-1 also exhibited a high degree of correlation along the rachis, although their relative intensities fluctuated (Figure 1C, C’, C’’). The Arp2/3 complex, represented by ARX-2, appeared in punctate structures along the rachis bridges, and CAP-1 shows strong co-localization with ARX-2 at the punctate structures (Figure 1C, C’, C’’). Thus, CAP-1 is an integral component of the actomyosin corset in the syncytial germline, associating with actomyosin as well as nucleators of both branched and linear F-actin.

### CAP-1 is required for maintenance of the germline architecture

Given the prominent localization of CAP-1 in the actomyosin corset, we sought to understand its role in the germline through loss of function experiments. We generated a *cap-1* null mutant by using CRISPR-Cas-9 to delete the entire *cap-1* gene. The homozygous null mutants, obtained from heterozygous mothers, successfully accomplish embryogenesis but arrest at the L3 larval stage. Since the homozygous *cap-1* mutants did not reach adulthood they were not useful for our study of CAP-1 in the germline and we therefore used RNAi-mediated knockdown (KD).

To assess the degree of KD in the germline, we carried out *cap-1* KD on worms expressing mKate2::CAP-1 and a membrane marker (GFP::PLC1δ-PH) (Supplementary figure 2A). We observed a ∼60% decrease in CAP-1 fluorescence in the *cap-1(RNAi)* germlines as compared to control worms (Supplementary figure 2B). Next, we subjected worms co-expressing the membrane marker and nuclear marker (HIS-58::mCherry) to *cap-1(RNAi)* starting from larval stage 1 (L1) and assessed their germline architecture in adulthood. We observed a range of phenotypes, which we classified broadly as ‘mild’ (53%) or ‘severe’ (47%). As shown in Figure 2A, CAP-1 loss of function led to multiple defects in gonad structure, including an abnormal loop region, abnormally sized oocytes (some multinucleated), defects in rachis tube morphology, and loss or collapse of germ-cell membranes. While in control germlines all the nuclei are held within the germ cells, we found nuclei floating in the rachis tube in the majority of *cap-1(RNAi)* gonads, both mild and severe (Figure 2B). Furthermore, while each germ cell has only one nucleus in the control, we found multinucleated cells in 9% ± 3% of mild and 27% ± 1.8% of severe *cap-1(RNAi)* germlines (Figure 2C).

**Figure 2:**
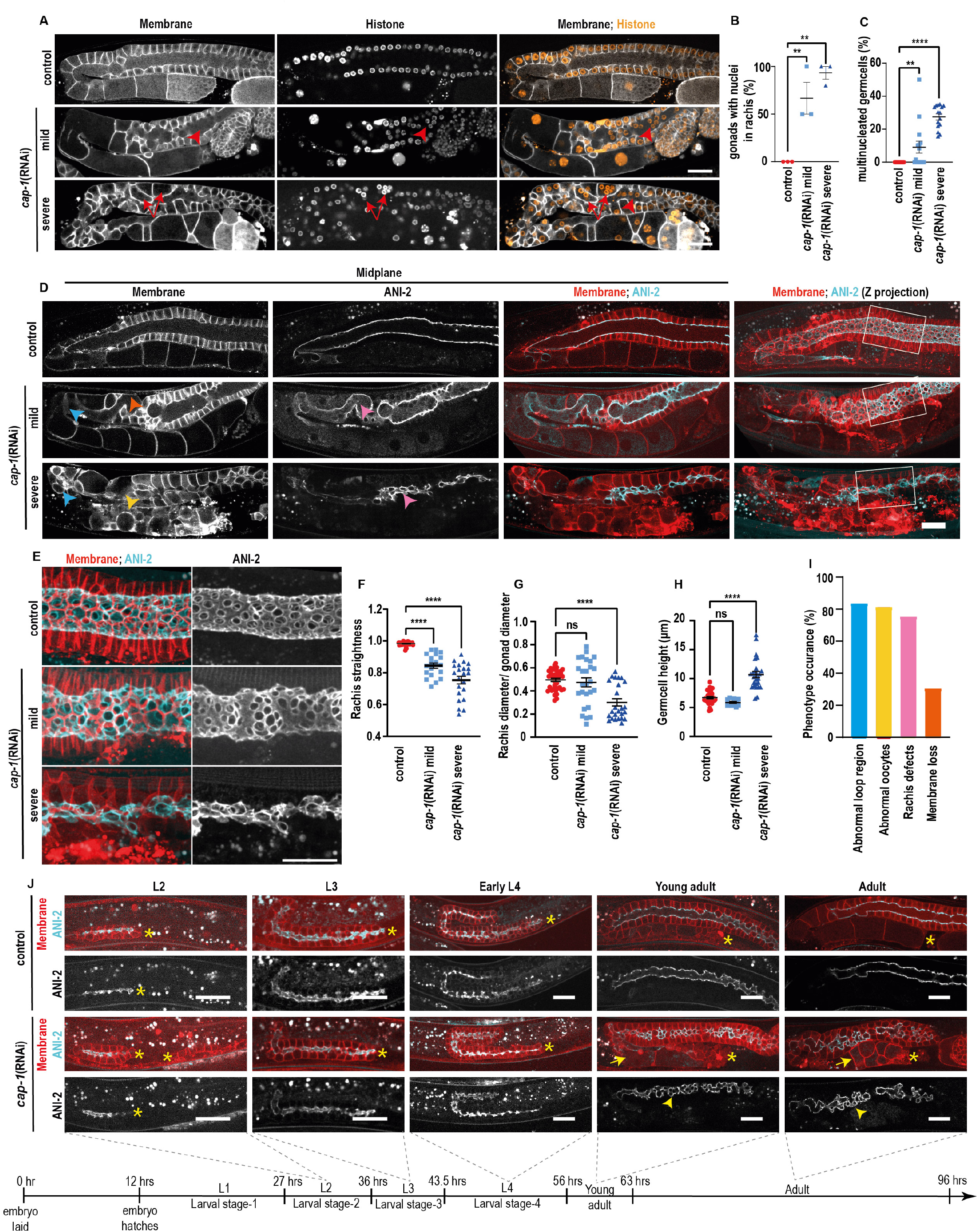
CAP-1 is required for the maintenance of germline architecture and functionality. (A) Confocal fluorescence images of the *C. elegans* germline expressing a membrane marker, GFP::PLC1δ-PH and a nuclear marker, HIS-58::mCherry. Top row shows control germlines while middle and bottom rows show *cap-1(RNAi)* germlines with mild and severe phenotypes. Arrowheads show nuclei mispositioned within the rachis. Arrows indicate multinucleated germ cells. (B, C) Percentage of gonads showing the phenotypes ‘nuclei in rachis’ and ‘multinucleated germ cells’ in gonads of control (N=32) versus mild (N=15) and severely affected (N=15) *cap-1(RNAi)* worms. (D) Confocal fluorescence images of the midplane and Z-projected views of gonads expressing a membrane marker, mCherry::PLC1δ-PH and a rachis bridge marker, ANI-2::GFP for control (top), mildly (middle) and severely (bottom) affected *cap-1(RNAi)* worms. Arrowheads indicate abnormal loop region (blue), abnormal oocytes (yellow), constricted and meandering rachis (pink) and membrane loss (orange). (E) Magnified views of the regions marked by white boxes in panel D, showing rachis morphology in gonads of control and *cap-1(RNAi)* worms. (F, G, H) Quantification of rachis straightness, rachis width and germ cell height in the gonads of control (N=38) versus mild (N=19) and strong *cap-1(RNAi)* (N=22) worms. (I) Percentage of phenotypes observed in CAP-1 KD gonads (N=48). (J) Confocal fluorescence images of germlines expressing the membrane marker (red) and ANI-2::GFP (cyan) from larval stage 2 till adulthood in control and *cap-1(RNAi)* worms. Asterisks indicate the proximal end of the germlines. Arrows and arrowheads indicate defective oocytes and rachis, respectively in *cap-1(RNAi)*. Scale bars, 20µm. Error bars are ± SEM. **p value < 0.01; ***p value < 0.001; ns, not significant (Student’s t-test).

To determine the effect of CAP-1 loss of function on rachis morphology with higher precision, we repeated the KD in a strain expressing a membrane marker (mCherry::PLC1δ-PH) and the anillin isoform ANI-2 tagged with GFP to mark the rachis bridges (Amini et. al. 2015). Unlike the rachis of control gonads, which was straight and maintained a fairly constant diameter, the rachis of *cap-1(RNAi)* worms was meandering and of variable diameter (Figure 2D, E). We quantified rachis straightness by dividing the length of a straight line between the two ends of the rachis by the length of a path along the centre of the rachis. In control worms, the rachis straightness was close to one, but it was significantly reduced upon *cap-1* KD (Figure 2F). Next, we measured the ratio between rachis and gonad diameters. In gonads mildly affected by *cap-1(RNAi)* the mean diameter ratio was not different from control, but the variance was higher. The severely affected gonads, on the other hand, exhibited a severely constricted rachis as compared with the control (Figure 2G). The decrease in rachis width is accompanied by an increase in germ cell height in the case of gonads severely affected by *cap-1RNAi* (Figure 2H). For an overview of the various phenotypes appearing in *cap-1(RNAi)* germlines we analysed 48 worms expressing a membrane marker and found that over 80% had abnormal oocytes and an abnormal loop region, over 70% had rachis defects, and ∼30% exhibited germ cell membrane loss (Figure 2I).

To determine whether the germline defects observed are due to *cap-1* depletion in the germline or whether *cap-1* depletion in other tissues might contribute to the phenotype, we performed *cap-1(RNAi)* in a germline-specific RNAi strain (DCL569), in which a null mutation in the RNAi-essential gene *rde-1* is rescued only in the germline by expression of *rde-1* under a germline-specific promoter (Zou et al., 2019). Following *cap-1(RNAi)* in this strain we observed the same germline defects as we found previously with whole worm RNAi and these defects appeared with the same penetrance and range of severities, suggesting that *cap-1* depletion within the germline is sufficient to generate these structural defects (Supplementary figure 2C).

To establish whether CAP-1 function is required during germline morphogenesis or for maintenance of its structure, we followed germline formation in *cap-1(RNAi)* conditions from the L2 stage until adulthood (Figure 2J). We did not observe any defect in gonad morphology during any of the larval stages. Starting from the young adult stage we could observe the onset of mild defects in gonad morphology in *cap-1* depleted worms, mostly oocyte morphology defects, and the defects increased in severity with time resulting in further gross structural defects in the adult germline. These results suggest that CAP-1 is not required during gonad development but is essential for maintenance of its structure in adulthood.

### CAP-1 depletion leads to increased F-actin and ARX-2 in the germline

To understand the dependence of germline structure on CAP-1 function we first set out to determine the effect of *cap-1(RNAi)* on the actin cytoskeleton. For this purpose, we chose to examine worms with a mild phenotype, because germline structure in the severely affected worms was too disrupted to compare with controls. F-actin, as visualized by phalloidin staining, is specifically enriched at rachis bridges in the germline, but is also observed at the basal and lateral sides of the germ cells, presumably marking their cortices. Phalloidin staining of dissected gonads showed a significant increase in F-actin in the germline of *cap-1(RNAi)* worms compared to control worms (Figure 3A). Quantification of fluorescence intensities showed that F-actin increased by approximately 2-fold at the rachis, lateral and basal side of the germ cells (Figure 3B-D). A similar increase was also observed with the endogenously GFP-tagged F-actin bundling protein PLST-1 in live worms (Supplementary figure 3A-D). Fluorescence recovery after photobleaching (FRAP) experiments did not reveal a significant difference in the turnover of PLST-1::GFP at the rachis bridges of *cap-1(RNAi)* worms, as compared to control worms (Supplementary figure 3E and 3G-I).

**Figure 3:**
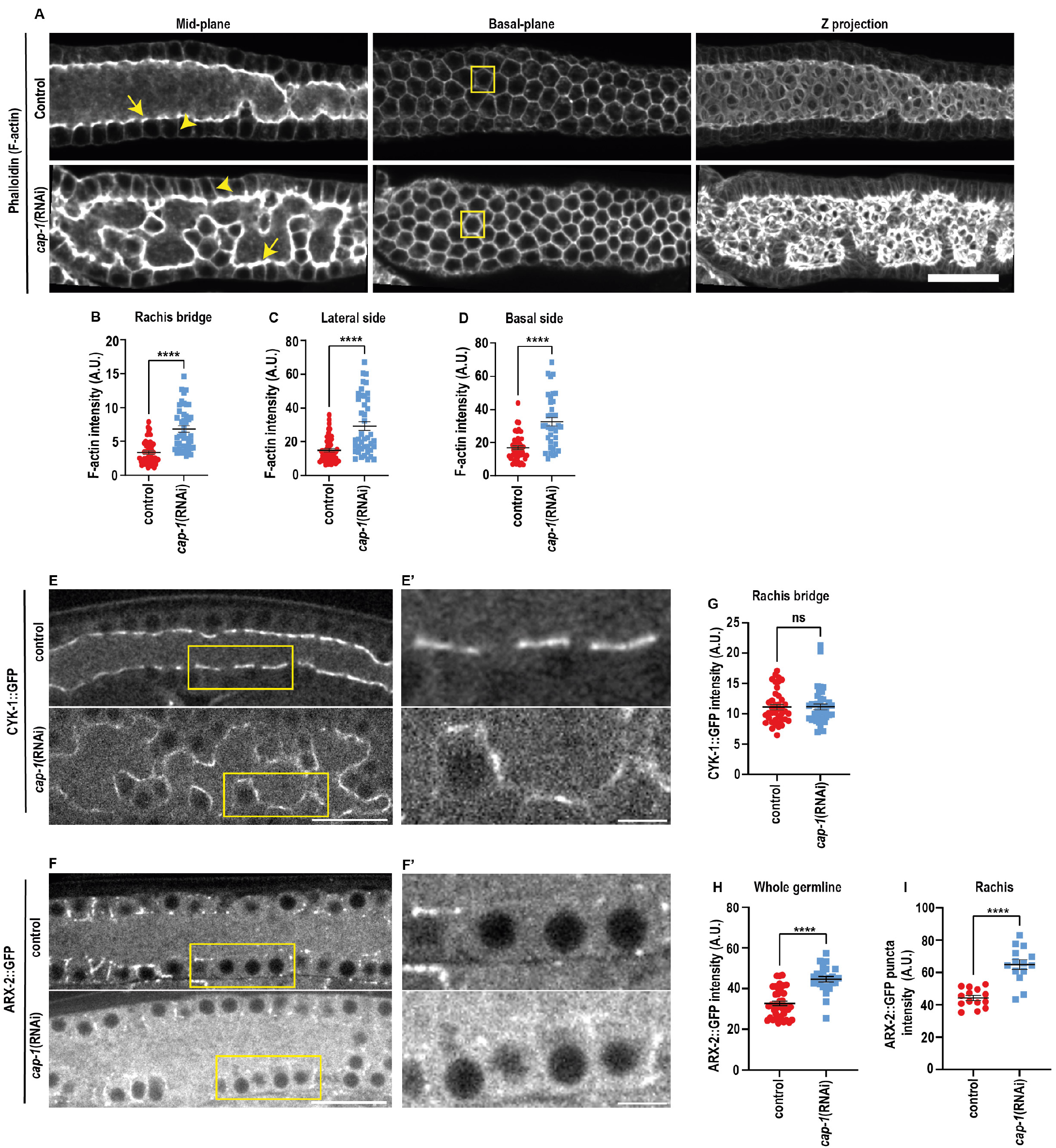
CAP-1 regulates F-actin and ARX-2::GFP levels in the *C. elegans* germline. (A) Midplane, basal plane and maximum-intensity Z-projected images of phalloidin stained germlines in control and *cap-1(RNAi)* worms. Scale bar, 20µm. (B, C, D) Quantification of mean F-actin intensity in untreated control (N=52) versus *cap-1(RNAi)* (N=42) at rachis bridges (arrows in A), lateral membranes (arrowheads in A) and basal sides of the germ cells (box in A). (E, F) Confocal fluorescence images of germlines expressing CYK-1::GFP and ARX-2::GFP in control versus *cap-1(RNAi)* worms. Scale bar, 20µm. (E’, F’) Magnified view of the rachis region marked by a yellow rectangle in panels E, F, respectively. Scale bar, 5µm. (G) Quantification of CYK-1::GFP intensity at the rachis bridges in control (N=41) versus *cap-1(RNAi)* (N=40) worms. (H, I) Quantification of ARX-2::GFP intensity in control versus *cap-1(RNAi)* worms within the whole germline (N=40, 25) and at the rachis (N=14, 14), respectively. Error bars are ± SEM. ***p value < 0.001; ns, not significant (Student’s t-test).

Given the substantial increase in F-actin levels in the germline, we explored whether *cap-1* KD affected the localization or levels of either of the known germline actin nucleators: the formin CYK-1 and the Arp2/3 complex. As shown earlier, CYK-1::GFP is enriched at the rachis, where it co-localized with CAP-1 (Figure 1E). We did not observe any difference in the localization or levels of CYK-1 in the germline following *cap-1(RNAi)* (Figure 3E, G). In contrast, we observed an average 1.3-fold increase in the levels of the Arp2/3 component ARX-2 in *cap-1(RNAi)* germlines (Figure 3F). The increase in ARX-2 was observed throughout the germline as well as within the punctate structures found along the rachis (Figure 3H, I).

### CAP-1 depletion leads to an increase in active NMY-2 at the germline corset

Given the increase in F-actin in the rachis following *cap-1* depletion and the known dependence of syncytial germline architecture on rachis actomyosin contractility, we investigated the effect of *cap-1(RNAi)* on non-muscle Myosin II/NMY-2 at the rachis. As shown in Figure 4A, we observed a 2-fold increase in NMY-2 levels at the rachis in *cap-1(RNAi)* (Figure 4A, B). Furthermore, using an antibody that recognizes only the active/phosphorylated form of Myosin II, we also observed an increase in the levels of phospho-myosin II at the rachis (Figure 4C, D), indicating an increase in contractility of the actomyosin corset in *cap-1(RNAi)* conditions. The observed increase in NMY-2 could, in theory, be due to more F-actin binding sites, to slower NMY-2 turnover, or a combination of both. In order to examine if the dynamics of NMY-2 were affected by *cap-1* depletion we carried out Fluorescence recovery after photo bleaching (FRAP) on endogenously tagged NMY-2::GFP at the rachis bridge. We did not observe a significant difference in the recovery kinetics of NMY-2 in *cap-1(RNAi)* versus the control, although we did observe a trend towards a larger immobile fraction in *cap-1* depleted background (Supplementary figure 3F, J, K, L). Thus, the increase in NMY-2 appears to be mainly due to the increase in F-actin.

**Figure 4:**
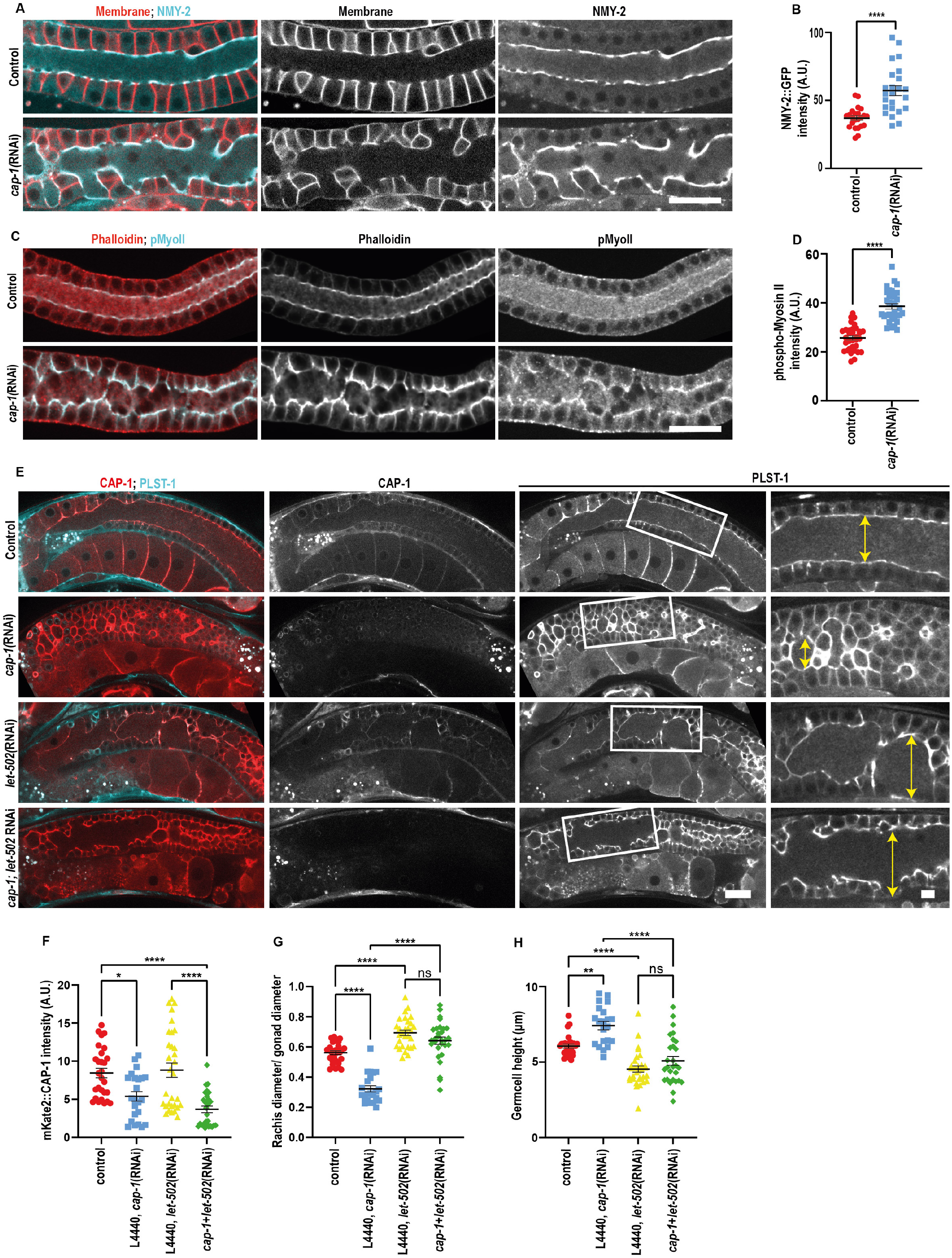
CAP-1 regulates contractility to maintain germline structure. (A) Confocal images of the germline expressing NMY-2::GFP (cyan) and GFP::PLC1δ-PH (red) in control and *cap-1(RNAi)* worms. Scale bar, 20µm. (B) Quantification of NMY-2::GFP intensity at the rachis bridge in control (N=23) versus *cap-1*(RNAi) (N=24) worms. Error bars represent ± SEM. ****p value <0.0001 (Student’s t-test). Scale bar, 20µm. (C) Confocal images of the germline in control and *cap-1*(*RNAi)* worms stained for phosphorylated Myosin II (pMyoII, cyan) and F-actin (phalloidin, red). Scale bar, 20µm. (D) Quantification of phospho-myosin II intensity at the rachis bridge in control (N=35) versus *cap-1(RNAi)* (N=31) worms. Error bars represent ± SEM. ****p value <0.0001 (Student’s t-test). Scale bar, 20µm (E) Midplane confocal views of the whole germline and magnified views of the rachis region (white box) in worms expressing PLST-1::GFP (cyan) and mKate2::CAP-1 (red) in control, *cap-1(RNAi), let-502(RNAi)* and *cap-1(RNAi);let-502(RNAi)*. Double-headed arrows mark rachis diameter. Scale bar, 20µm, scale bar in magnified images, 5µm. (F, G, H) Quantification of mKate2::CAP-1 intensity, rachis width and germ cell height in germlines of control (N=28), *cap-1(RNAi)* (N=23), *let-502(RNAi)* (N=31) and *cap-1(RNAi);let-502(RNAi)* (N=29) worms. Error bars are ± SEM. *p value < 0.1; **p value < 0.01; ****p value <0.0001 (One-way ANOVA test).

### Increased actomyosin contractility can explain the germline defects of CAP-1 loss of function

The observed increase in F-actin and active NMY-2 at the rachis led us to hypothesize that *cap-1* loss of function phenotypes is due to increased contractility of the rachis actomyosin corset. To test this idea, we first compared germline morphology of *mel-11(RNAi)* with *cap-1(RNAi)*. MEL-11 is the *C. elegans* myosin phosphatase, and its depletion is known to lead to an increase in actomyosin contractility (Piekny and Mains, 2002). Importantly, we observed similar morphological changes, namely a constricted rachis and increased germ cell height in *cap-1(RNAi)* and *mel-11(RNAi)* (Supplementary figure 3M). If indeed *cap-1* germline defects are due to increased rachis contractility, then reducing myosin activity should rescue these defects. We tested this idea by depleting the myosin activator Rho kinase/LET-502 in *cap-1(RNAi)* worms. As evident in figure 4E, *let-502(RNAi)* displayed a phenotype opposite of that of *mel-11(RNAi)* or *cap-1(RNAi)*, namely it had a wider than normal rachis and shorter germ cell membranes. We performed the double RNAi KD in worms expressing mKate2::CAP-1 and quantified CAP-1 intensity to confirm *cap-1* depletion and rule out a dilution effect (Figure 4E and 4F). Germlines with the double KD of *let-502* and *cap-1* resembled the *let-502 (RNAi)* phenotype of a broad rachis and short germ cell membranes, indicating that the morphological defects observed in *cap-1(RNAi)* gonads are a result of increased contractility of the actomyosin corset in the syncytial germline (Figure 4G, H).

## Discussion

Tissue morphogenesis involves coordination between multiple cells undergoing structural and identity changes and generating pushing and pulling forces against each other to ultimately assume a functional shape (Heer and Martin, 2017). Tissue homeostasis is also dependent on a fine balance between external and internal mechanical forces. Here, we found actin CP CAP-1 to be a critical component of the *C. elegans* germline rachis actomyosin corset, where it co-localizes with F-actin as well as formin and Arp2/3 and regulates its contractility.

Different actin structures varying in architecture, connectivity and contractility can exist within the same cell or tissue and accomplish different functions in a spatiotemporally regulated manner. For example, the fission yeast cytoskeleton has four distinct actin structures, which differ largely from each other in structure, function and mechanical properties (Kovar et al., 2011). CP associates with actin patches in the fission yeast to exclude formin from the Arp2/3 complex-mediated actin structures, hence preserving the identity of distinct actin structures (Billault-Chaumartin and Martin, 2019). We found CAP-1 in the syncytial *C. elegans* germline to be enriched at rachis bridges, which differ from the rest of the germ cell cortices in terms of composition and function (Amini et al., 2014; Priti et al., 2018). The formin, CYK-1 exhibits an enriched localisation at the rachis bridge while the Arp2/3 component ARX-2 does not, suggesting a differential distribution of branched and linear actin networks in the germline. However, we did not observe spreading of formin into Arp2/3 domains in the absence of CAP-1. Instead, we observed an increase in F-actin and, to a lesser degree, an increase in ARX-2 levels everywhere in the germline. The increase in F-actin can be explained by the absence of a factor stopping its polymerization. The mechanism responsible for increasing ARX-2 is not clear and may be due to an as-of-now uncharacterized transcriptional response or simply by the availability of more F-actin. Nevertheless, since the increase in F-actin exceeded the increase in ARX-2 levels, the density of branched actin in the network likely decreased, altering the balance between linear and branched actin in favour of linear actin. We therefore propose that loss of CAP-1 results in a change in the architecture of the actin network at the rachis bridge, which supports a common role for CAP-1 in generating and maintaining distinctly different actin structures in a cell or tissue.

Actin and non-muscle myosin II combine to form contractile structures, which are regulated spatiotemporally by several accessory proteins. Multiple actin binding proteins, including filamin, fascin, anillin, ezrin, and moesin, have been shown to affect actomyosin contractility via regulation of different aspects of actin organisation (Murrell et al., 2015). We found a dramatic increase in NMY-2 levels at the rachis upon CAP-1 KD, a result that was confirmed by phospho-myosin II staining. Our FRAP experiments did not detect a significant change in NMY-2 dynamics following CAP-1 KD. On the other hand, the twofold increase in NMY-2 levels at the rachis closely matches the twofold increase in F-actin following CAP-1 KD. We therefore conclude that the increase in myosin levels is primarily due to the availability of more F-actin.

The balance between linear and branched actin can also influence the contractility of the system as a whole. *In-vitro* reconstitution assays and modelling have shown that the balance between linear and branched actin networks influences the level of ‘network connectivity’, which can affect myosin mediated contractility according to a bell-shaped curve (Belmonte et al., 2017; Ennomani et al., 2016). We do not know where along this connectivity curve the rachis actomyosin network is in wild-type conditions, but it is possible that a decrease in connectivity due to the decreased density of Arp2/3 branching results in increased contractility. Alternatively, the increase in contractility upon *cap-1* depletion might be despite the decrease in connectivity and is solely due to the increase in linear F-actin and NMY-2.

The consequences for the germline of increased rachis contractility due to CAP-1 loss of function are disastrous for both the structure and function of the tissue. While development of the germline proceeds normally in the absence of CAP-1, maintenance of the syncytial germline structure requires CAP-1 and in its absence multiple structural defects appear. The increased contractility results in a highly constricted rachis, accompanied by defective oocytes. Importantly, inhibiting actomyosin contractility in the CAP-1 KD condition, by inhibiting myosin phosphorylation by Rho Kinase, is able to rescue the restricted rachis phenotype, strengthening our conclusion that CAP-1 affects rachis morphology through actomyosin contractility. Actin CP has been shown to affect actin organisation in *Drosophila* bristles and oocytes (Gates et al., 2009; Hopmann and Miller, 2003). Studies in cells have highlighted the importance of CP in cytoskeletal organisation and actin-driven protrusive activity (Fujiwara et al., 2014; Johnston et al., 2018; Jung et al., 2016). To the best of our knowledge, our study is the first to demonstrate a role for CP in regulating actomyosin contractility in a tissue. We found CAP-1 to be strongly expressed in multiple tissues in the adult worm and in the embryo. Future studies will reveal the role CP plays in other tissues and establish which are tissue-specific and which are general functions of CP *in vivo*.

## Acknowledgements

This study was supported by Israel Science Foundation grant 767/20 awarded to RZB, and scholarships from the Mechanobiology Institute Singapore and the Israeli Council for higher education awarded to S.R. Some strains were provided by the CGC, which is funded by NIH Office of Research Infrastructure Programs (P40 OD010440). We thank Edwin Munro and David Kovar (The University of Chicago) for illuminating discussions.

## Author contributions

Conceptualization, S.R., P.A. and R.Z.B; Investigation, S.R. and P.A.; Formal Analysis, S.R. and P. A.; Visualization, S.R.; Writing – Original Draft, S.R; Writing – Review & Editing, P.A. and R.Z.B.; Supervision, R.Z.B.; Funding Acquisition, R.Z.B.

## Materials and Methods

### Worm strain maintenance

All *C. elegans* strains were grown and maintained on NGM (nematode growth medium) plates seeded with OP50 bacteria according to standard protocols (Brenner, 1974). All worms were kept at 20ºC, unless specified otherwise. All strains used in this study have been listed in the table. Some strains in this study have been obtained from the CGC (University of Minnesota) and some have been made by InVivo Biosystems (Eugene, OR, USA) using CRISPR-Cas9 knock-in technology.

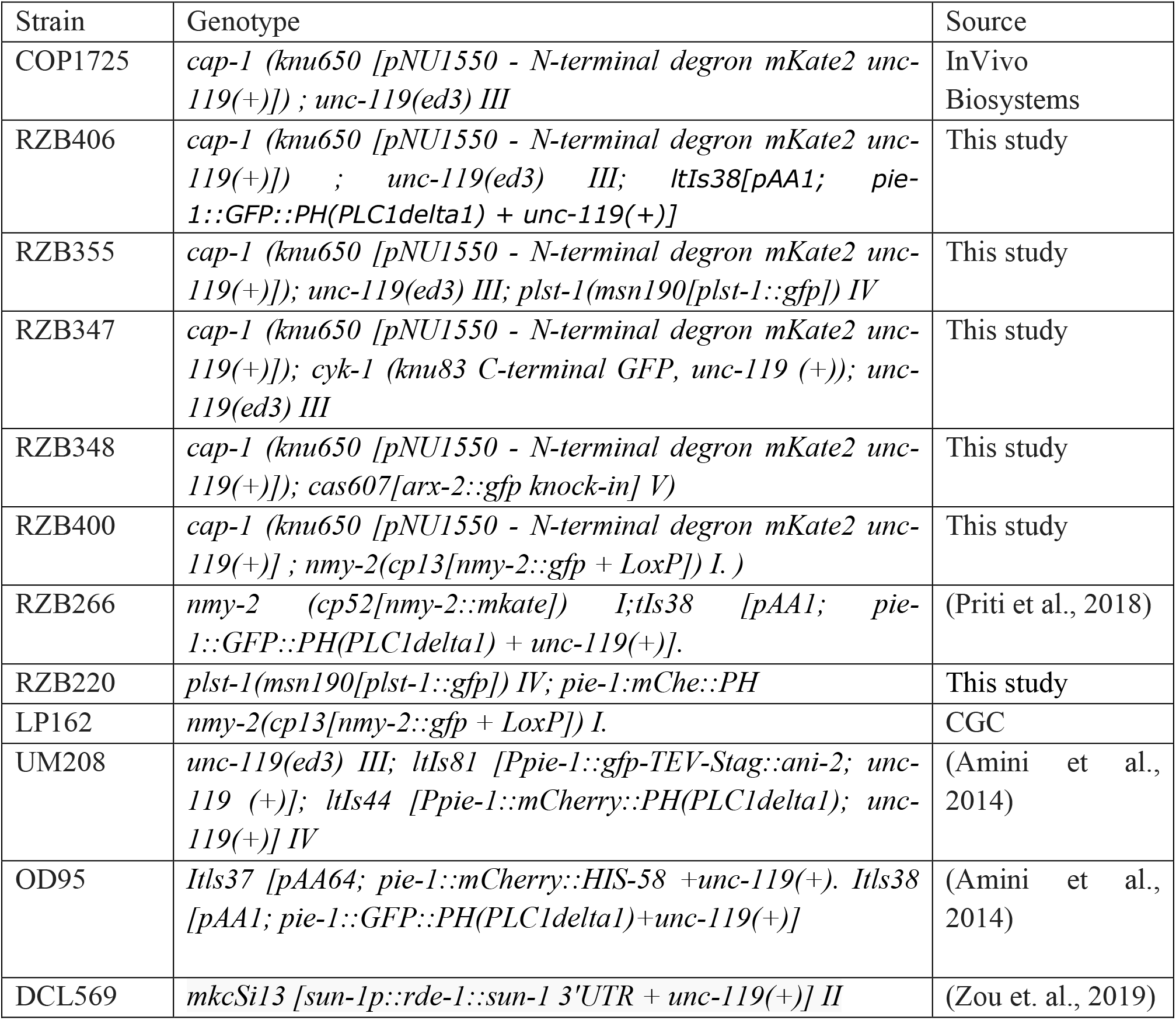

### RNA interference

All RNAi experiments were performed by feeding RNAi. Bacterial clones of HT115(DE3) bacterial strain expressing the vector L4440 containing gene specific sequences were obtained from the Ahringer or Vidal libraries (Source BioScience) and sequenced for confirmation. RNAi plates were prepared with NGM media containing 1mM IPTG and 100 μg/ml of ampicillin. A bacterial clone containing L4440 vector alone was used as a negative control for the experiments, unless specified otherwise. For dsRNA induction, an overnight culture of bacteria was diluted 1:50 and grown at 37ºC with shaking for 3-4 hours, after which IPTG was added to a final concentration of 1mM and allowed to grow for an additional 3-4 hrs. This culture was used to seed the RNAi plates, air dried and grown overnight at room temperature. Gravid hermaphrodites were bleached on the RNAi plates to release the embryos (unless specified otherwise). Worms hatched on the RNAi plate were grown to adulthood, when they were collected either for immunofluorescence or live imaging. UM208 mild phenotype exhibiting worms were put on the RNAi plates at L3 stage and grown till adulthood.

### Phalloidin staining

Gonads were dissected in M9 buffer containing 0.5mM levamisole on a glass slide. The dissected gonads were fixed in 3.5% formaldehyde in PBS for 20 minutes, then permeabilised in PBS with 0.025% Triton-X100 for 5 minutes. Next, the gonads were incubated in phalloidin-TRITC (Sigma, P1951) in the dark at room temperature. The gonads were washed with PBS to remove excess phalloidin and were mounted on a glass slide with vectashield mounting medium (Vector lab, Cat. No. H-1000) and mounted for imaging.

### Immunofluorescence

Dissected gonads (in 0.5mM levamisole in M9 buffer) were washed to remove levamisole and fixed with 3.5% formaldehyde in PBS for 20 minutes. Next, the gonads were washed thrice in PBS and then permeabilised with 0.25% tween in PBS for 10 mins. The gonads were then washed thrice with PBS, incubated in blocking solution (1% BSA, 0.1% Tween and 30mM glycine in PBS) at room temperature for 1 hour. A 1:400 dilution of anti phosphoMLC Ser19 (Cell Signaling Technology, Cat. no. 3671) in blocking solution was used to incubate the gonads at 4ºC overnight. After 3 washes with PBS, the gonads were incubated with a solution containing 1:500 dilution of anti-rabbit secondary antibody conjugated with Alexa 488 (Invitrogen, Cat. no. A21244), 1:250 phalloidin-TRITC (Sigma, Cat. no. P1951) and 1:1000 DAPI (Sigma-Aldrich) in blocking buffer at room temperature for 1.5 hours. After 3 washes with PBS, the gonads were stored in vectashield (Vector lab, Cat. no. H-1000) and mounted for imaging.

### Microscopy and image acquisition

Live adult hermaphrodites in 10mM levamisole or immunostained dissected gonads stored in vectashield were mounted on fresh 3% agarose pads prepared on a glass slide. Imaging was carried out with a Nikon Ti2E microscope equipped with a Yokogawa W1 spinning disk system and a Plan Apo 60X oil 1.4 NA and a Plan Apo 100X oil 1.45 NA. Samples were illuminated with 405nm, 488nm or 561nm lasers (Gataca systems, France) for excitation and acquired on a prime 95B sCMOS camera (Photometrics, Tucson, AZ). The software Metamorph version 7.10.2.240 (Molecular Devices, CA) was used as the controlling interface. All images were captured with Z stacks of 1μm spacing.

### Image analysis

All image analysis was carried out with the help of FIJI (Schindelin et al., 2012). For measurement of germ cell height, several germ cells were selected in the pachytene region in the Z plane exhibiting the germline midplane. Germ cell height was measured for selected cells. The ratio of germline width/rachis width was measured at three regions in the germline midplane and the average ratio was calculated for each gonad.

Line scan intensity plot: to analyse the relative localisation between mKate2::CAP-1 and PLST-1::GFP, NMY-2::GFP, ARX-2::GFP or CYK-1::GFP at the rachis, a 5 pixel thick line was drawn along the rachis and the fluorescence profile was generated for both channels with Fiji and graphs were plotted with GraphPad Prism 9.

Fluorescence intensity measurement: each image was corrected for the background fluorescence before measurement. The mid plane of the germline rachis was selected for analysis of PLST-1::GFP, CYK-1::GFP, NMY-2::mKate and phalloidin at the rachis bridges using Fiji. Mean intensity was measured along a 5-pixel wide line drawn along the rachis bridges for both control and RNAi conditions. For measurement of PLST-1::GFP and phalloidin at the basal and lateral plane, the basal and lateral plane of the germline was selected respectively. The segmented line tool was used to trace the cell boundaries at the basal plane and the cell junctions at the lateral plane and mean intensity was calculated.

Total ARX-2::GFP fluorescence was calculated by tracing the germline outline at the midplane and mean intensity was measured after background subtraction. For measurement of ARX-2 puncta fluorescence, the rachis midplane was outlined and thresholded for selecting the puncta and the mean intensity was measured.

#### Fluorescence recovery after photobleaching (FRAP)

FRAP experiments were performed with the iLAS2 module (Gataca systems, Massy, France) for targeted laser illumination on the spinning disk microscope described above with a Plan Apo 100X oil 1.4 NA objective. A circular region of interest along the germline rachis of worms expressing NMY-2::GFP was selected manually and photobleached with the 488nm laser at 80-100% laser power. Images with five Z slices (1μm spacing) were acquired using 50% 488nm laser and exposure time of 400ms at an interval of 30secs for a total duration of 10 mins. The average intensity of the ROI was measured with Fiji for the bleached, unbleached and background ROIs. The fluorescence intensity of the bleached region was double normalised for photobleaching and background using the double normalisation method. The final analysis of difference in recovery rates was done with Graphpad Prism 9 as performed in (Priti et al., 2018).

### Statistical analysis

All statistical analysis was carried out with the software Graphpad Prism 9. Student’s t-test or one-way analysis of variance (ANOVA) was used to carry out tests of significance for two samples or more than two samples, respectively. All graphs are represented as mean ±S.E. The sample number for each experiment is indicated in the corresponding figure legend.

