## Supplementary file for "Actin capping protein regulates actomyosin contractility to maintain germline architecture in *C. elegans*"

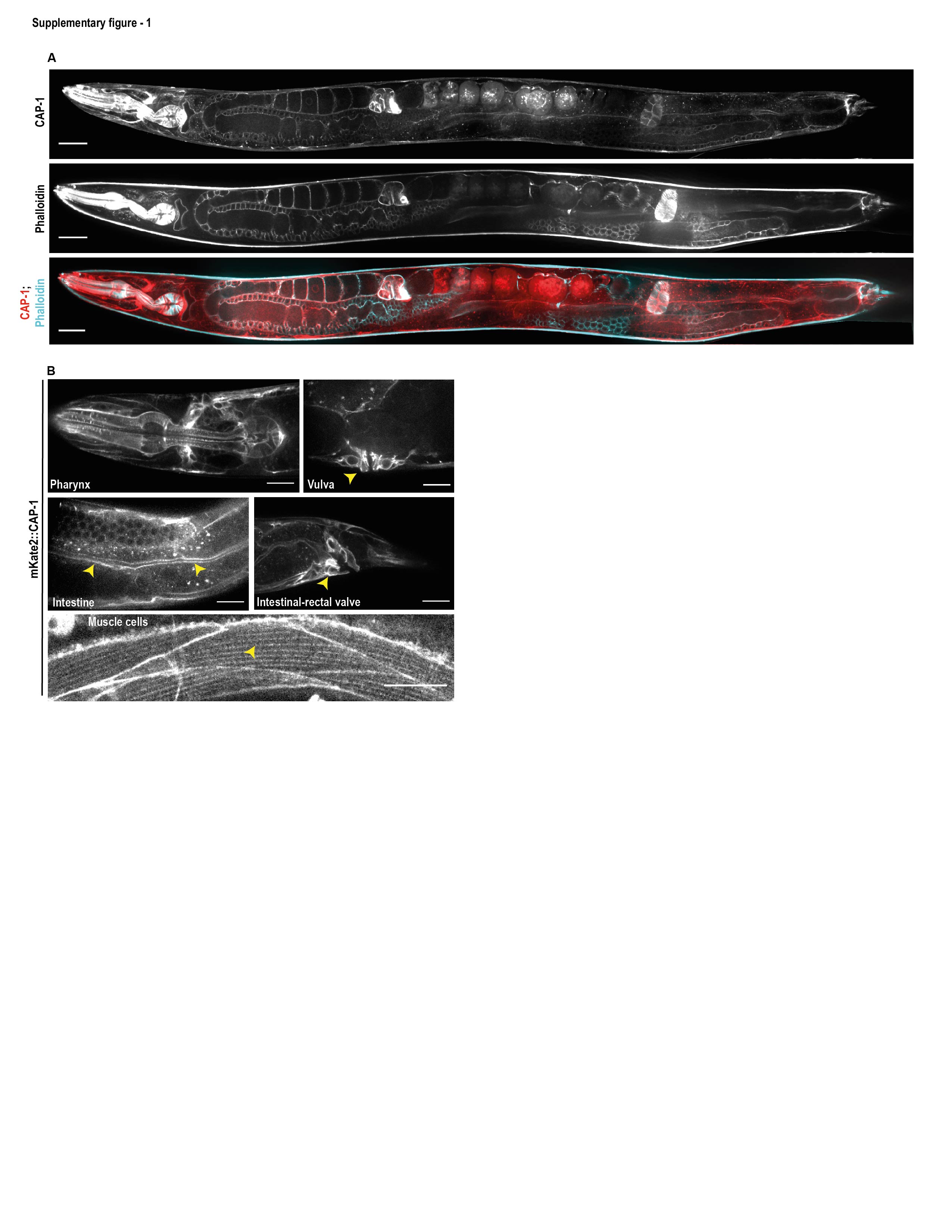

**Supplementary Figure 1: Localization of endogenously mKate2 tagged CAP-1 in multiple tissues.**

(A) Representative confocal fluorescence image of a phalloidin stained (cyan) *C. elegans* hermaphrodite expressing mKate2::CAP-1 (red).

(B) Confocal fluorescence images of the localization of mKate2::CAP-1 in the pharynx, vulva, intestine, rectal valve and muscle striations. Arrowheads indicate CAP-1 localisation in respective tissues. All scale bars, 20 µm.

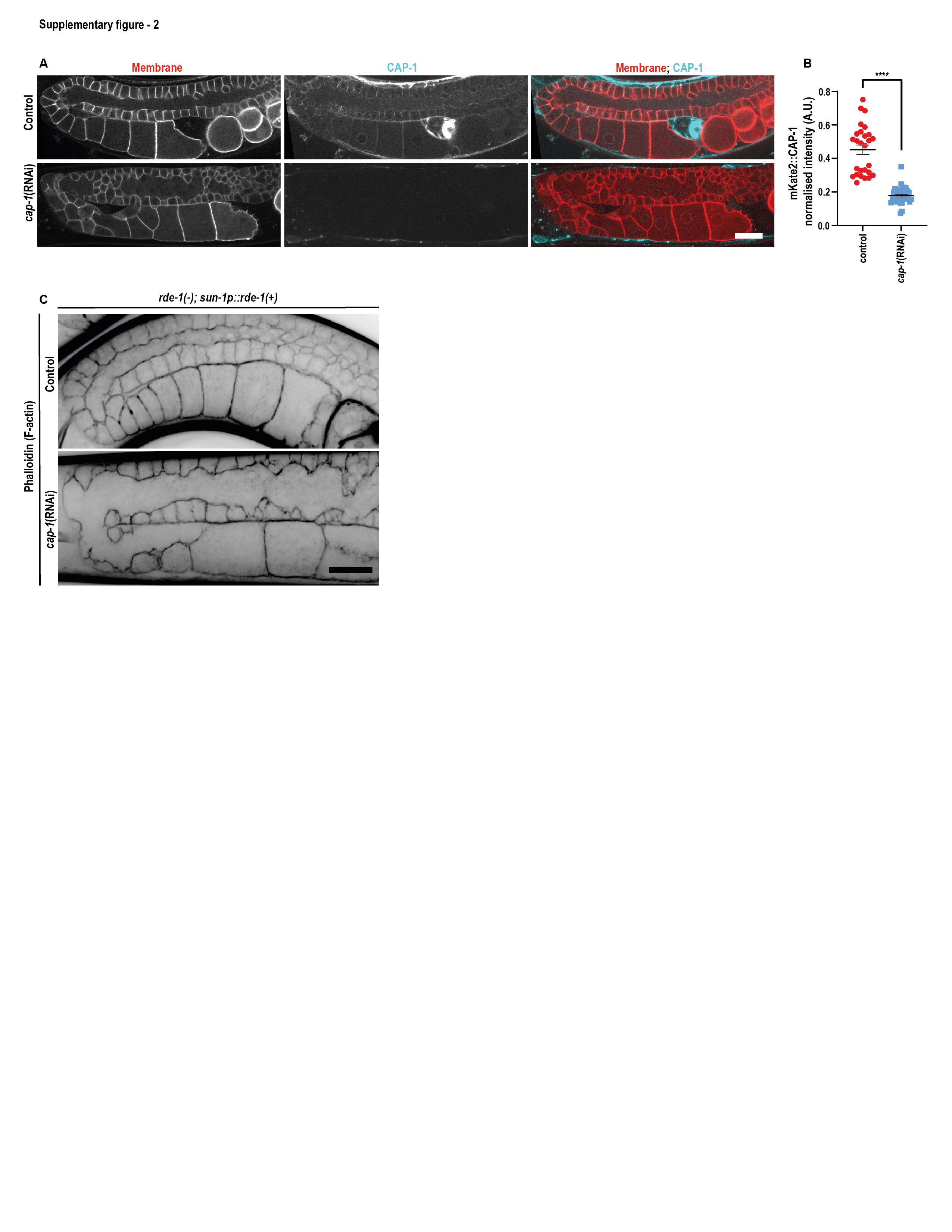

**Supplementary Figure 2: ­CAP-1 is required tissue-autonomously to regulate germline architecture**

(A) Confocal fluorescence images the *C. elegans* germline expressing GFP::PLC1δ-PH (red) and mKate2::CAP-1 (cyan) in control and *cap-1(RNAi)* worms.

(B) Quantification of normalized mean mKate2::CAP-1 intensity in control(N=27) versus *cap-1(RNAi)* (N=40) worms at the rachis bridge. Error bars are ± SEM. ***p value < 0.001 (Student’s t-test).

(C) Phalloidin stained images of the *C. elegans* germline deficient in somatic tissue RNAi (*rde-1*(-); *sun-1p::rde-1*(+)) in control and *cap-1(RNAi)* worms.

Scale bars, 20 µm

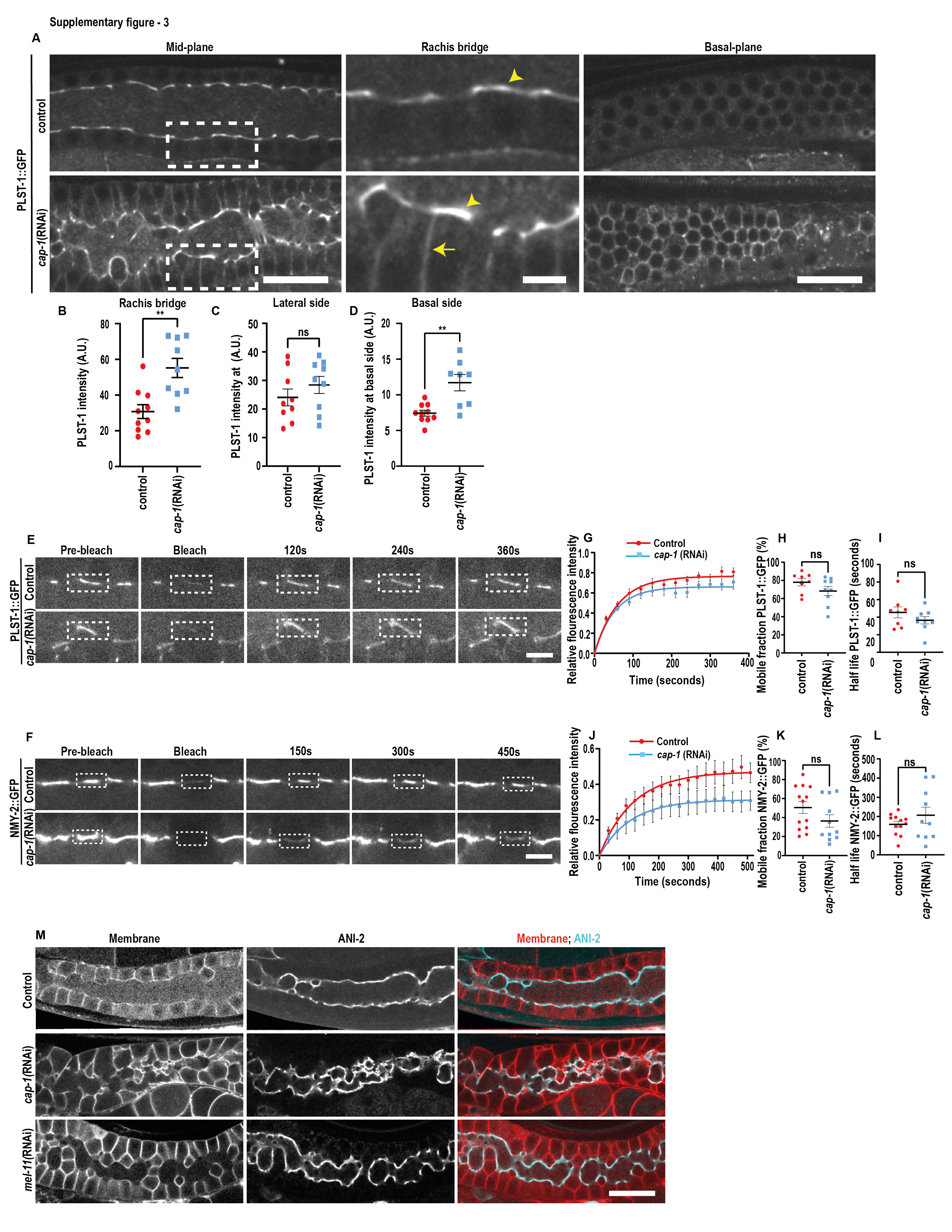

**Supplementary Figure 3: CAP-1 regulates PLST-1::GFP level but does not significantly affect the turnover of PLST-1::GFP and NMY-2::GFP**

(A) PLST-1::GFP in control and *cap-1(RNAi)* gonads. Left column shows mid-plane view; middle column consists of magnified views of the regions marked by white rectangles in the left column, showing increased intensity of PLST-1::GFP at rachis bridge (arrow) and lateral germ cell membrane (arrowhead). Right column shows basal views of the same germline. Scale bar in left and right columns, 20 µm; scale bar in magnified images, 5 µm.

(B, C, D) Quantification of mean intensity for PLST-1::GFP at rachis bridge, lateral side and basal side of the germ cell membrane in control (N=9) and *cap-1(RNAi)* (N=10) worms.

(E, F) Time-lapse confocal images of PLST-1::GFP showing the fluorescence recovery of PLST-1::GFP and NMY-2, respectively, after photo bleaching at the rachis bridge in control and *cap-1(RNAi)* worms. Scale bar, 5 µm.

(G-L) Quantification of recovery kinetics, mobile fraction, and half-life of PLST-1::GFP (N=8, 8) and NMY-2::GFP (N=11, 13), respectively, in control and *cap-1(RNAi)* worms.

(M) Representative confocal fluorescence images of the *C. elegans* germline expressing membrane marker, mCherry::PLC1δ-PH (red) and rachis bridge marker, ANI-2::GFP (cyan) in untreated control, *cap-1(RNAi)* and *mel-11(RNAi)* worms. Scale bar, 20 µm.

Error bars are ± SEM. **p value < 0.01; ns, non-significant (Student’s t-test).
